# Calling differential DNA methylation at cell-type resolution: avoiding misconceptions and promoting best practices

**DOI:** 10.1101/2021.02.28.433245

**Authors:** Han Jing, Shijie C. Zheng, Charles E. Breeze, Stephan Beck, Andrew E. Teschendorff

## Abstract

The accurate detection of cell-type specific DNA methylation alterations in the context of general epigenome studies is an important task to improve our understanding of epigenomics in disease development. Although a number of statistical algorithms designed to address this problem have emerged, the task remains challenging. Here we show that a recent commentary by Rahmani et al, that aims to address misconceptions and best practices in the field, continues to suffer from critical misconceptions in how statistical algorithms should be compared and evaluated. In addition, we report contradictory results on real EWAS datasets.

---

While a growing number of Epigenome-Wide Association Studies (EWAS) have identified DNA methylation (DNAm) changes associated with a wide range of phenotypes and exposures^1,2^, the biological interpretation of these molecular alterations remains challenging due to the underlying cell-type heterogeneity of the tissues used in such studies. One solution would be to perform EWAS in purified cell-types, but this remains costly and has mostly only been carried out in immune cells ^3–6^. Thus, there is an ongoing attempt to address this challenge computationally, by devising statistical algorithms that can detect cell-type specific differential DNAm, i.e. differentially methylated cell-types (DMCTs) ^7,8 9–11^, from EWAS performed in bulk-tissue.

Most of the proposed statistical algorithms fall into a class of conceptual models where the DNAm data, represented by say “X”, is modelled in terms of how the phenotype/exposure, represented by say “Y”, affects DNAm, which can broadly speaking be denoted statistically as (X|Y) (read X given Y). These class of (X|Y) models are sensible for EWAS studies that aim to understand how an exposure or phenotype such as smoking or obesity may affect DNAm patterns in specific cell-types ^7^. One of these algorithms, called CellDMC ^7^, explicitly models the DNAm data in this way, naturally incorporating statistical interaction terms between the phenotype/exposure and the estimated cell-type fractions in a linear modelling framework. This model is based on the intuition that differential methylation differences in a given cell-type would be larger in cases for which the affected cell-type is more abundant.

An entirely different conceptual model and algorithm called TCA was proposed by Rahmani and colleagues^9^. Importantly, the acronym TCA stands for Tensor Composition Analysis, reflecting an attempt by the authors to infer a full data tensor defined over features (CpGs), samples and cell-types, from which subsequent association analyses can then be performed in each cell-type separately. We agree with Rahmani et al that TCA is a more expressive model than CellDMC, but disagree that this by itself “automatically leads to better performance”, specially on real EWAS data. Indeed, models with more parameters are more prone to overfitting, a key elementary concept in machine learning ^12^ that Rahmani et al seem unaware of. In their TCA paper, Rahmani and colleagues propose a number of different implementations of TCA, of which only one truly captures the spirit of TCA. This is the implementation where one infers the underlying tensor. However, this implementation is not the recommended one by the authors. Instead, the authors go on to present another version of their “TCA” method, which like CellDMC directly infers DMCTs, but which in contrast to CellDMC/TOAST, models the phenotype/exposure (Y) in terms of the DNAm data (X), which in their recent commentary they denote as “TCA (Y|X)”. This particular version of TCA can come in three different flavours, depending on the statistical test that one can apply: (i) a joint-test to test a combined effect over all cell-types, (ii) a marginal conditional test where the marginal test for a given cell-type is conditioned on all other cell-types, and (iii) a marginal-test where one tests for each cell-type separately and independently from the other cell-types.

In their original TCA paper, Rahmani and colleagues performed a direct comparison of TCA (Y|X) to CellDMC, concluding that TCA (Y|X) outperforms CellDMC. However, as we argued at great length in a recent critical review ^13^, the analysis and results of the TCA paper were incomplete and biased, rendering the conclusions of the original TCA paper invalid. In their recent reply, Rahmani and colleagues claim “misconceptions”, yet as we argue further below, the new data presented by Rahmani and colleagues fully support and justify our original criticism. Moreover, Rahmani et al’s commentary continues to suffer from key critical misconceptions, which we elaborate on further below, raising further major concerns regarding their “recommended best practices”.

Our original criticism of the TCA paper ^9^ was based on the following facts:

1. The authors presented the TCA (Y|X) model, which was compared to CellDMC (a X|Y model) using evaluation metrics that were not designed to test DMCT-calling methods, and which therefore led to a biased comparison. Indeed, the definition of sensitivity as used in the TCA paper naturally favors a method implementing a joint or marginal test, as implemented in TCA (Y|X) but not a method implementing a marginal conditional test, as implemented in CellDMC. Thus, <u>the very same origin of the misconception that Rahmani et al accuse other authors of, is present in their own TCA paper and in the main figures of their paper</u>.
2. Methods were never compared in terms of the precision metric, where precision, also known as positive predictive value (PPV), is related to the false discovery rate (FDR) by PPV=1-FDR, a key metric that assesses the fraction of true positives among significant associations. When comparing DMCT-calling methods to each other, it is critical to consider such a metric, yet Rahmani et al ignored this metric for unknown reasons. In the context of the CellDMC paper, the PPV metric was not critical, since the CellDMC paper did not compare CellDMC to any other DMCT-calling method, only to DMC-calling methods, as indeed there were no other DMCT-calling methods to compare to.
3. Methods were never compared in terms of their computational efficiency. Only now, in their recent commentary, do Rahmani et al acknowledge that this was a major omission of their TCA paper.
4. The authors ignored publicly available matched DNAm and FACS-sorted based cell-count data in blood as a benchmark for assessing how accurate TCA (Y|X) can estimate cell-type fractions.
5. The authors only assessed their TCA (Y|X) method on one real EWAS dataset in Rheumatoid Arthritis (RA), for which the existence of a gold-standard list of associations is highly questionable, with the only such available list ^5^ being completely ignored, again for unknown reasons. In our CellDMC paper, we assessed CellDMC in 4 different real EWAS scenarios, including breast cancer, endometrial cancer, a smoking EWAS in buccal swabs and RA. In the case of RA, we used the validated list of RA-associated CpGs in B-cells ^5^. Given the large dimensionality of omic datasets and the danger of overfitting to random variation in data, the validation of a method in only one real data context is not a recommended practice ^12^.

In their recent commentary ^14^, Rahmani et al effectively acknowledge most of our criticism and also follow our recommendations, adding the PPV metric to their evaluations, and including smoking EWAS, a scenario for which a more reasonably gold-standard set of loci exists ^15^. Most importantly however, they also alter their method, presenting a new TCA (X|Y) model. Yet, in the abstract of their commentary, they state that we “misused” their TCA method, falsely claiming that the TCA (X|Y) model was “part of their original TCA paper”. In our critique of the TCA paper, we implemented TCA using the recommended settings as specified in their TCA R-package. Specifically, the TCA method implemented in their TCA R-package (version-1.1.0) was TCA (Y|X), with the recommended marginal model as the default, which is the one we therefore used, since this is also the version of TCA that was used in their original TCA paper, and for which comparative results to CellDMC were presented in the main figures of their TCA paper. Thus, <u>there was no misconception on our side</u>: we were bound to use the same implementation and flavor of TCA that were used in the original TCA paper when directly comparing to CellDMC. Thus, the origin of any misconceptions can be traced back to the main figures of their own TCA paper where they compare an (Y|X) model to an (X|Y) model (CellDMC), a comparison which by the author’s same arguments (as given in their recent commentary) is not meaningful.

A related misconception by Rahmani et al is that they compared the TCA (Y|X) joint and marginal models to our CellDMC algorithm, which by construction implements a marginal conditional test, using only the sensitivity metric for evaluation, ignoring the all-important PPV metric. Had Rahmani et al included the PPV metric in their evaluations, this would have alerted them to the very low precision of the TCA (Y|X) model, as demonstrated by us ^13^. Indeed, it is precisely because we pointed out to them the need to consider the PPV metric, that Rahmani et al have now altered their recommended TCA method from “TCA (Y|X) marginal” to “TCA (X|Y) marginal conditional”, which we note is now a very similar model to CellDMC, since CellDMC is also an (X|Y) model implementing a marginal conditional test. This explains why in Fig.1 of their recent reply, TCA (X|Y) and CellDMC perform so similarly.

**Fig.1:**
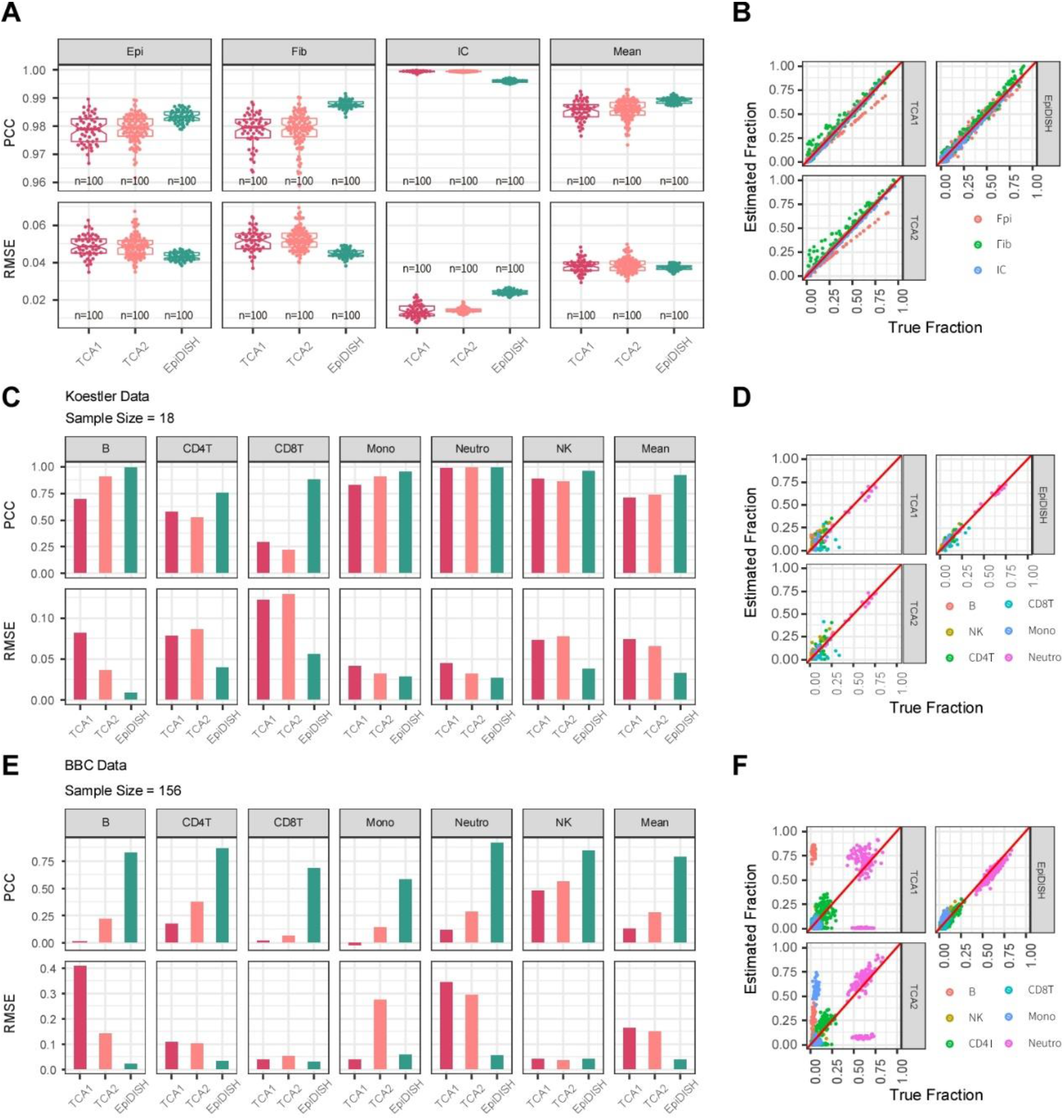
Evaluation of cell-type fraction estimates using TCA. **A)** Boxplots of the Pearson Correlation Coefficient (PCC) and Root Mean Square Error (RMSE) between the estimated cell-type fraction and the true fraction for 3 cell-types (Epi=epithelial, Fib=fibroblast, IC=immune cell), for the case of 100 in-silico generated mixtures of epithelial (Epi), fibroblast (Fib) and immune cell (IC) subtypes. The last panel shows the mean PCC and RMSE over the 3 cell-types. Shown are the PCC and RMSE for 3 different methods of estimating the cell-type fractions: TCA1=using final TCA estimates initializing with EpiDISH, TCA2=using final TCA estimates initializing with EpiDISH plus noise, EpiDISH=cell-type fraction estimates derived using the robust partial correlation framework of EpiDISH ^18^. **B)** Corresponding scatterplots of estimated vs. true cell-type fractions for the 3 cell-types and for each of the 3 different methods. **C-D)** As A-B), but now comparing the estimated cell-type fractions to the true fractions in 18 whole blood samples from Koestler et al ^17^. In this dataset, the true fractions refer to either FACS cell counts or to known proportions for the experimentally derived mixtures, as described in Koestler et al ^17^. **E-F)** As C-D), but now for a whole blood data set of 162 samples with matched FACS cell counts.

The above argument highlights another key and much more serious misconception made by Rahmani et al, one that does not only appear in their original TCA paper, but one that recurs again in their recent commentary. Indeed, <u>it is critically important to note that when we evaluate a DMCT-calling method on simulated data, that we should use exactly the same method in all different simulation scenarios</u>. This is critically important because if simulation is to provide objective guidance on how a method will perform on real data, then we should not be allowed to use prior insight from the simulation set-up to tailor the choice of method to the given simulation scenario. For instance, in the Methods subsection of their TCA paper, titled “Power Simulation”, the authors state “ ...*We considered three scenarios across a range of effect sizes as follows: different effect sizes for different cell types (using a joint test),…and a scenario with only a single associated cell type (a marginal test)*.” In other words, the authors use a different implementation of their TCA (Y|X) method (joint-test or marginal-test) depending on the simulation scenario (all cell-types changing or only one cell-type changing). This is tantamount to using a different method for different simulation scenarios, and would not lead to a meaningful assessment, because <u>in a real EWAS situation we would never know in advance if a CpG is altered in one cell-type or in multiple cell-types</u>. Given that in a real EWAS situation, all potential combinations of altered cell-types may be realized, it seems therefore impractical to propose application of all potential methodological variations, as Rahmani et al are proposing in their recent reply, and especially so, if the underlying method is computationally demanding or even prohibitive. Indeed, the original TCA paper, as well as their recent commentary, lack clarity as to which specific implementation of TCA should be used in practice, and most importantly how to synthesize the results from the application of all various implementations of TCA. In contrast, CellDMC only implements a marginal-conditional test, and exactly this and only this method was evaluated across all simulation scenarios and all 4 real EWAS datasets in Zheng et al ^7^. Thus, CellDMC could be applied after an ordinary DMC-calling method to ensure high sensitivity and precision, a computationally efficient solution which Rahmani et al once again miss to mention.

In addition to these misconceptions by Rahmani et al, several of our earlier major concerns, as described by us previously ^13^, remain unanswered:

First, we had argued that an important evaluation metric is the estimation of cell-type fractions themselves. Indeed, the TCA method involves an iterative procedure, in which initial cell-type fractions are re-estimated ^9^. A natural question to address therefore is whether the final cell-type fraction estimates agree better with the ground truth values, a question which is perfectly addressable not only within a simulation framework, but also in the context of real blood EWAS for which matched FACS blood cell counts are available. We assessed this using both simulation and real blood DNAm datasets with matched FACS cell counts, concluding that TCA’s final cell-type fraction estimates are significantly worse than the initial ones (**Fig.1**). This contradicts the result shown in SuppFig.2 of the TCA paper, where in our opinion the authors used the wrong benchmark. For whole blood, reliable DNAm references exist ^16–18^ and yet the authors of the TCA paper initialize their TCA method with relatively poor cell-type fraction estimates, leading to the illusion that the final TCA estimates are good enough. Moreover, the authors of the TCA paper offer no assessment on real DNAm data with matched flow-cytometric cell counts, despite such data being available ^19^. Our analysis on both simulated and real DNAm data demonstrates that TCA’s final cell-type fraction estimates are worse than those obtained using a DNAm reference (**Fig.1**).

Second, when comparing algorithms a key evaluation metric is computational efficiency, yet the original TCA-paper did not compare methods in terms of their computational efficiency. This is particularly pertinent when the task at hand is computationally demanding. Because TCA infers a data tensor where the genomic feature (CpG) dimension is fairly large, it is a computationally very demanding task. In contrast, CellDMC scales linearly with the number of CpGs and does not take much longer to run than a standard linear regression model. Indeed, we performed a detailed comparison of the two methods in terms of their runtimes, finding that CellDMC is much more efficient (**Fig.2**). For instance, running TCA on a typical EWAS study would take a day or more on a standard workstation, whilst CellDMC would complete the task in about 10 to 20 minutes (**Fig.2**), which represents an approximate 100-fold improvement over TCA. To illustrate this further, we applied TCA to an in-house EWAS dataset consisting of 3500 samples all generated on EPIC methylation beadarrays (~850,000 CpGs). We ran TCA on a powerful Dell PowerEdge R815 server with 512 GB RAM and 50 cores, yet after running for 2 full days, TCA crashed due to large memory requirements. In contrast, CellDMC completed the same task on the same server and same number of cores in just under 10 minutes. In their recent commentary, Rahmani et al claim 10-fold improvements in computational speed, but refrain from testing their method on larger EWAS encompassing thousands of samples. This plainly contradicts their own recommendation to favor TCA over CellDMC in EWAS containing more than 60 samples.

**Fig.2:**
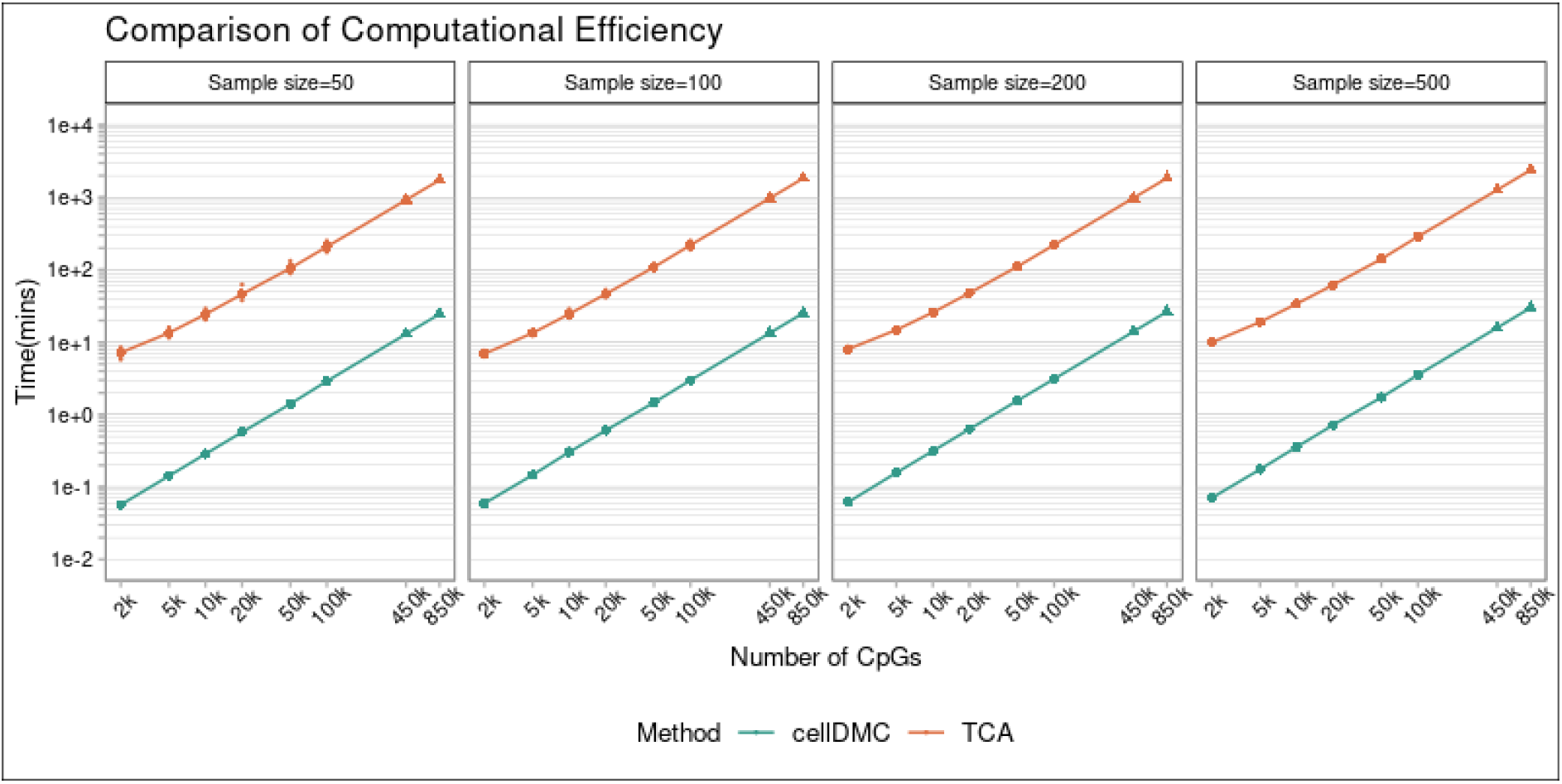
Runtimes of CellDMC and TCA. Runtimes (in minutes, shown in a log-base-10 scale) of CellDMC and TCA for 4 different sample sizes and different numbers of CpGs (x-axis, also shown on a log-base-10 scale). Runtimes were obtained on a Dell PowerEdge R815 server running CentOS 7.7.1908. Total RAM was 512 GB and while this server has 64 cores, the estimates in this table were obtained using only 1 core (to avoid potential confounding by server-load, which was in any case low when running these processes).

Third, evaluation on simulated data can serve to highlight which methods may work better on real data, but there is no guarantee that this would be the case. Thus, evaluation on real EWAS datasets is of key importance. In this regard, we and others have argued that there are two key conditions that real data must satisfy. First, methods need to be assessed in relation to phenotypes where there is some biological ground truth, or at least, where there is strong evidence and experimental validation for the implication of specific differentially methylated positions in relation to the phenotype. Second, multiple independent EWAS cohorts is required in order to ensure that results obtained in one dataset generalize to others ^19^. In the TCA paper, the authors compare TCA to CellDMC only in the context of one Rheumatoid Arthritis (RA) EWAS study ^20^, whilst CellDMC was evaluated in the context of four different real EWAS datasets and phenotypes (smoking, RA, breast and endometrial cancer) ^7^. In our opinion, sole reliance on RA to evaluate performance of a method on real data should not be used as a conclusive argument, because there is little experimental validation data to support the definition of a gold-standard list of RA-associated CpG sites, not least in specific blood cell subtypes. In fact, one of the few studies which did validate RA-associated loci in blood cell subtypes (specifically in B-cells) and across multiple cohorts is the one by Julia et al ^5^, yet Rahamani et al chose to ignore it for unclear reasons. We previously used this data to validate CellDMC’s application on the same RA EWAS study considered in the TCA paper, concluding that CellDMC did have the sensitivity to detect these experimentally validated RA-associated DMCTs ^7^.

In any case, and as demonstrated by us recently ^15^, a better phenotype to consider would be smoking, because by now there is a well-established and highly reproducible gold-standard list of smoking-associated DMCs derived from whole blood EWAS ^21^, with follow-up studies assessing these loci in specific blood cell subtypes ^22,23^. In Su et al ^22^, a list of 7 CpGs were identified, 5 of which exhibit myeloid specific smoking-associated DNAm changes, with 2 exhibiting lymphoid-specific changes. We thus compared CellDMC to TCA across 3 large independent EWAS cohorts with available smoking information, two in whole blood ^20,24^ and one in buccal swabs ^25^ (which contain about 50% immune-cells ^26^), to explore what these methods predict for the gold-standard list of 7-CpGs. Whereas CellDMC’s predictions were largely in agreement with those from Su et al and also largely consistent between the 3 cohorts, TCA was unable to predict the specificity of the myeloid and lymphoid specific smoking DMCTs (**Fig.3**).

**Fig.3:**
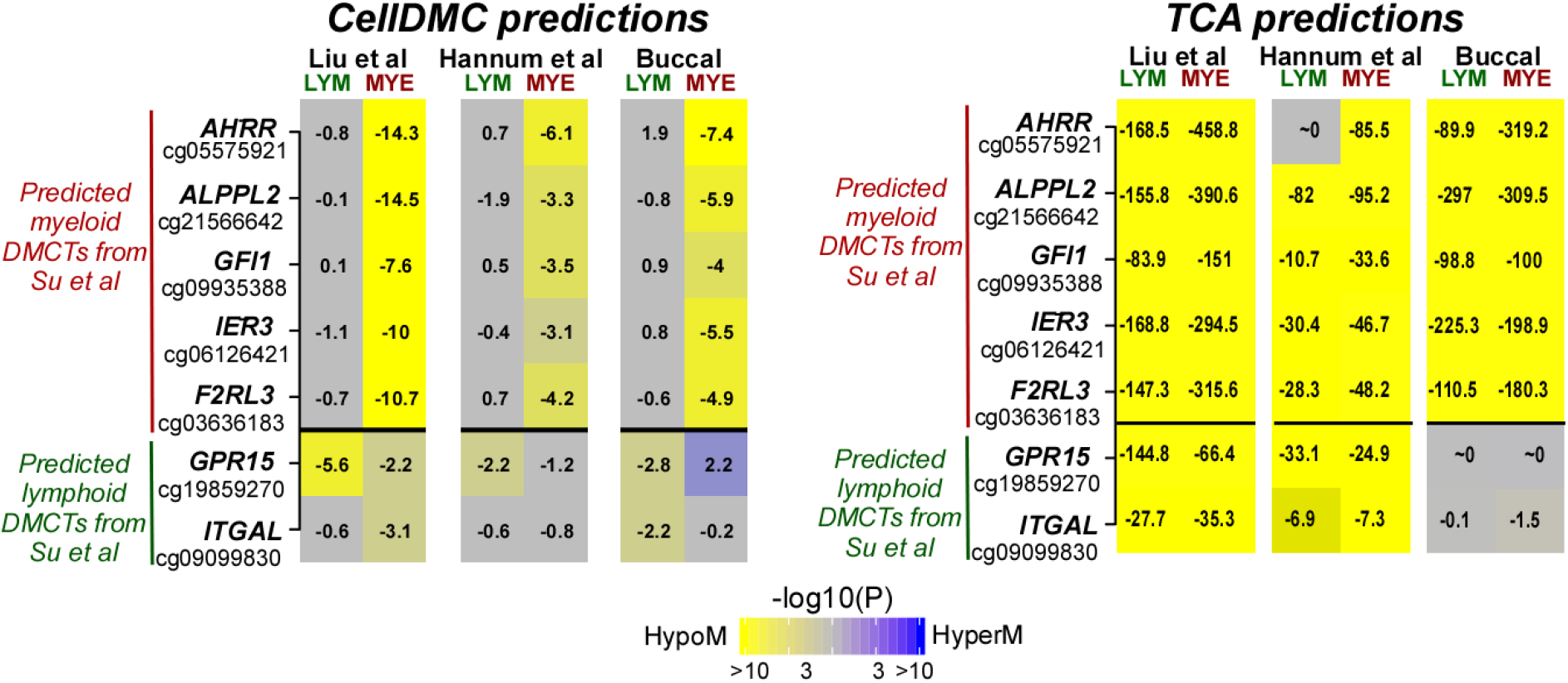
Comparison of CellDMC to TCA in three smoking EWAS. For 7 CpGs which Su et al identified via experimental validation to be exhibiting myeloid and lymphoid specific DNA hypomethylation in blood of smokers, we display the estimated statistics of association for CellDMC and TCA, for each of these 7 CpGs across 3 independent cohorts (two in whole blood and one in buccal swabs, which contains about 50% immune cells). The estimated statistics are displayed as heatmaps of statistical significance with yellow indicating strong hypomethylation and blue strong hypermethylation. The numerical entries are the corresponding t-statistics (CellDMC) or LRT-statistics (TCA) from which the P-values are derived. Statistics and P-values are shown for the myeloid (neutrophil+monocyte) and lymphoid lineages (T-cells, B-cells and NK-cells). TCA was run with the tcareg function, as recommended by the authors, with the same confounders as included in CellDMC.

Importantly, we note that in their recent commentary, Rahmani and colleagues present, oddly enough, the TCA results for a joint-test, which clearly leads to wrong predictions, alongside the results for TCA being implemented in “tensor-mode” using the *tca* function, which by definition and construction falls into the (X|Y) models. However, according to our data, results for implementing TCA in this tensor mode are as poor as implementing it in the (Y|X) marginal mode (**Fig.4**), which therefore contradicts their recent findings.

**Fig.4:**
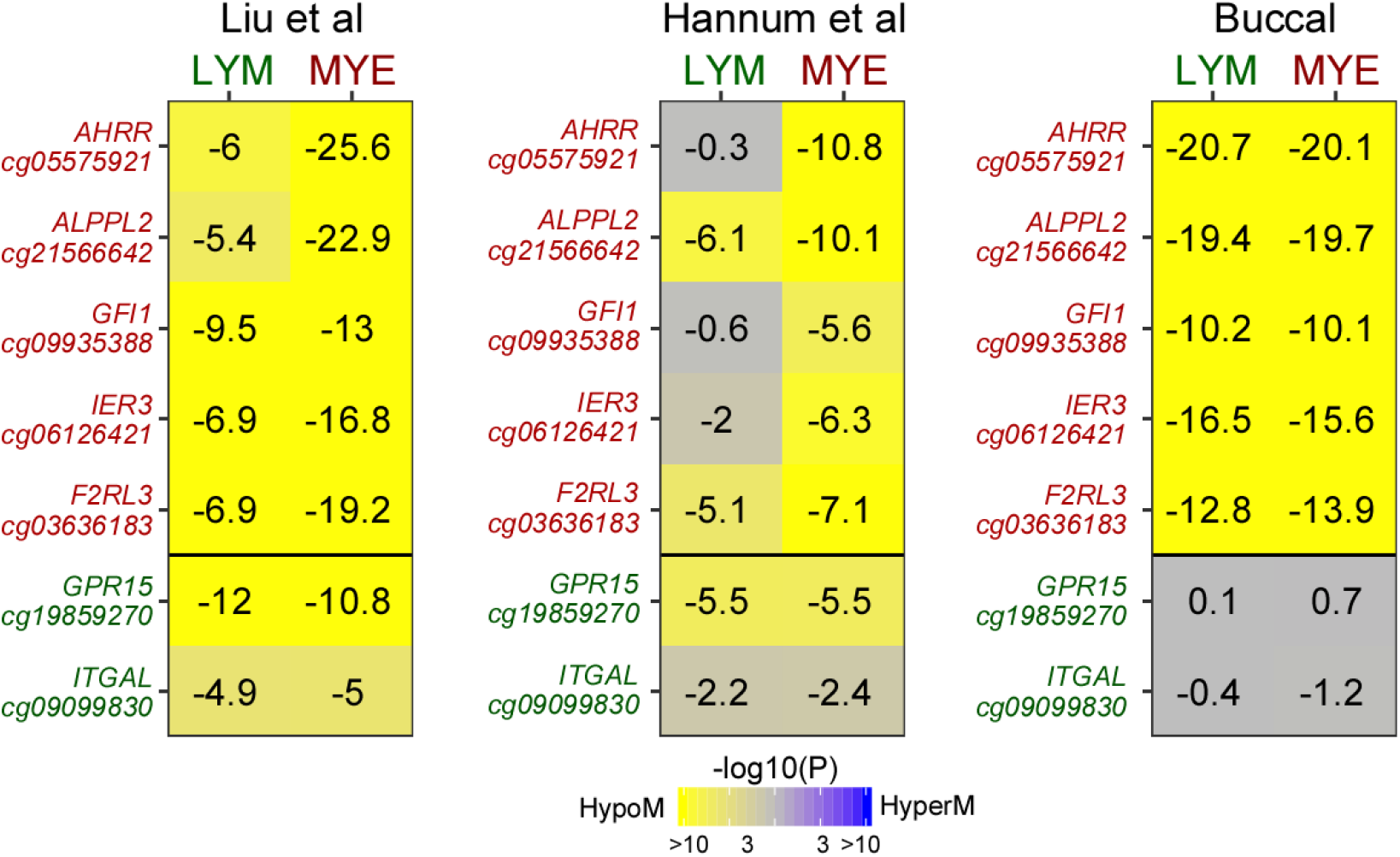
Evaluation of TCA in tensor-mode in three independent smoking EWAS. For 7 CpGs which Su et al identified via experimental validation to be exhibiting myeloid (top-5, shown in red) and lymphoid (bottom-2, shown in green) specific DNA hypomethylation in blood of smokers, we display the estimated t-statistics of associations of TCA (TCA run in tensor-mode) for these 7 CpGs across 3 independent cohorts (two in whole blood and one in buccal swabs, which contains about 50% immune cells). The estimates are displayed as heatmaps of statistical significance with yellow indicating strong hypomethylation and blue strong hypermethylation. The numerical entries are the corresponding t-statistics from which the P-values are derived. Statistics and P-values are shown for the myeloid (neutrophil+monocyte) and lymphoid lineages (T-cells, B-cells and NK-cells). We note that here TCA was run, first inferring the tensor, and then running separate regressions for each cell-type, whereas in Fig.3 we used the tcareg function.

Moreover, Rahmani et al claim that CellDMC exhibits a low precision and specificity in these smoking EWAS. We can easily refute this claim by drawing attention to the community to our recent meta-analysis study of smoking EWAS ^15^, where CellDMC displays excellent specificity and precision, as evaluated on over 7 independent smoking EWAS studies, and further supported by results obtained on realistic simulated data. Thus, our results on real EWAS contradict those presented by Rahmani et al.

Reproducibility is critical given the complexities of applying these algorithms to simulated as well as real data. Further details of the methodological implementations are provided in **Supplementary Methods** document, and we have also submitted our R-scripts and data, for generating the results on both simulated as well as real data, to the figshare repository: *https://doi.org/10.6084/m9.figshare.12922322*

In summary, identification of DMCTs at high sensitivity and precision is imperative to maximize the value of EWAS. Our previous objective assessment ^13^, as well as updated analyses presented here demonstrate that overall, CellDMC achieves reasonably high sensitivity, specificity and precision over a wider range of different biological scenarios, and at a much lower computational cost compared to TCA. Indeed, while the new TCA (X|Y) model proposed by Rahmani et al is a substantial improvement over their TCA (Y|X) algorithm, there is no clear evidence of an improvement of TCA (X|Y) over CellDMC on simulated or on real EWAS data.

## Declarations

### Author Contributions

Statistical and computational analyses were done by JH with input from SZ, and separately also by AET. CEB and SB contributed valuable feedback. Study was conceived and designed by AET. Manuscript was written by AET.

### Ethics approval and consent to participate

Not applicable, as this study only analyses existing publicly available data.

