## Supplementary Methods for "Calling differential DNA methylation at cell-type resolution: avoiding misconceptions and promoting best practices"

*Simulation model:* Our simulation models are based on the Illumina 450k DNA methylation (DNAm) profiles for 6 major immune cell subtypes (CD4+ and CD8+ T-cells, Granulocytes, monocytes, CD19+ B-cells and natural killer (NK) cells) from Reinius et al [1]. In this DNAm set there are 6 purified samples for each cell-type. In order to more realistically simulate real mixtures of these 6 cell-types in large numbers of samples, we first estimated means and standard deviations at each CpG site across each of the 6 cell types assuming an underlying beta-distribution using the method of moments. For generating the in-silico mixtures and for computational efficiency, we constructed the DNAm data matrices over a total of 1333 CpGs, made up of 1000 CpGs picked at random from the 450k array and the additional set of 333 "reference" CpGs which make our EpiDISH DNAm reference matrix [2]. These 333 reference CpGs are used to infer cell-type fractions in mixtures. A total of 200 in-silico mixtures were generated, i.e. 100 controls and 100 cases where DMCTs are defined. To define differentially methylated cell-types (DMCTs), that is differentially methylated cytosines within specific cell-types, we first selected at random 100 CpGs from the 1000 non-reference CpGs, but ensuring that these 100 CpGs exhibit either low (<0.2) or high (>0.8) average DNAm levels in the specific cell-types to be altered. This was done to ensure that we could consider a wide-range of effect sizes without having to worry about the bounded nature of beta-values. Thus, these levels of methylation represent the baseline levels in control samples, and thus for CpGs exhibiting low methylation in controls we would only consider hypermethylation in cases, and vice-versa for CpGs highly methylated in controls, we would only consider hypomethylation. In total, we considered 4 different DMCT scenarios: unidirectional change in one cell type, unidirectional changes in two cell types, bidirectional changes in two cell types and bidirectional changes in three cell types. Thus, in the unidirectional two cell-type scenario, a given CpG that is normally unmethylated in say granulocytes and B-cells, would exhibit hypermethylation in both cell-types in cases. Similarly, in the bidirectional two cell-type scenario, a CpG that is normally unmethylated in granulocytes and fully methylated in B-cells, would exhibit hyper and hypomethylation in granulocytes and B-cells, respectively. For each scenario, we considered a total of 60 different effect sizes with values ranging from 0 to approximately 12, with corresponding absolute DNA methylation differences in individual cell types (between cases and controls) varying from 0 to 0.6 maximally. Thus, these range of effect sizes capture effectively any realistic scenario. The calculation of effect size we use is given by the following equation:

$$effect\_size = \frac{\Delta\mu}{\sqrt{\frac{\sigma_1^2 + \sigma_2^2}{2}}}$$

where $\Delta\mu = |\mu_1 - \mu_2|$ is the absolute mean DNA methylation shift between case and control group, $\mu_1$ and $\mu_2$ are the mean DNA methylation levels in case and control group respectively, and where $\sigma_1$ and $\sigma_2$ are the variance of DNA methylation levels in case and control groups, respectively. Here we fixed $\sigma_1 = \sqrt{\frac{\mu_1(1-\mu_1)}{\mu_2(1-\mu_2)}} \cdot \sigma_2$ to ensure that the variance changes with the mean according to a beta-valued distribution.

Cell-type fractions to use for each in-silico mixture were obtained by sampling from a true immune cell subtype fraction distribution, as derived using EpiDISH on the Hannum et al whole blood dataset [3], which contains 656 Illumina 450k whole blood samples. Finally, we composed bulk methylation level at each CpG site by taking a linear combination of the simulated cell-type-specific levels across the 6 major immune cell subtypes, weighted by the corresponding cell type fractions. For every scenario and effect size, we ran 10 Monte-Carlo simulations. This was a sufficient number per effect-size, given that we considered a fairly large number (60) of different effect sizes.

*Implementation of CellDMC and TCA on simulated data:* We implemented CellDMC [4] using the corresponding R package named "EpiDISH". CellDMC was run with the estimated cell type fractions from EpiDISH, which effectively corresponds to the true fractions, as shown by us previously [2, 4]. Case/control status was encoded as a binary phenotype. Correction for multiple-testing was done by estimating the FDR and calling significance at an FDR threshold of 0.05, which results in an output matrix containing predicted DMCT(s) per cell-type, as well as their direction of DNAm change.

We applied TCA using the corresponding R package called "TCA" [5]. We ran TCA with initial cell-type fraction estimates as given by EpiDISH, that is we use the same estimates as when running CellDMC. In our simulation model, there are no other sources of variation besides phenotype and cell-type, so the parameters C1 and C2 were set to NULL. From the TCA-fit, we subsequently inferred the data-tensor defined over cell-types, CpGs and samples. Thus, for each cell-type, this data tensor defined a corresponding cell-type specific DNAm data matrix, on which we subsequently performed an ordinary linear regression to derive t-statistics and P-values of association against case/control status. Multiple-testing correction was performed as before using an FDR < 0.05 threshold, resulted in an output matrix containing predicted DMCT(s) per cell-type, as well as their directionality of DNAm change. We note that we also applied TCA using the *tcareg* function, since the authors claim that this demonstrates improved sensitivity over the 2-step procedure of first inferring the tensor and subsequently running separate regressions for each cell-type. However, all

TCA results reported in this work are independent of the specific implementation.

*Definition of sensitivity (SE), specificity (SP) and positive predictive value (PPV):* All these measures were first defined for each cell-type separately, and subsequently, values were averaged over all relevant cell-types. For a given cell-type that was chosen to be altered, the sensitivity to detect DMCTs in this cell-type was defined as the ratio of the predicted true number of DMCTs divided by the total number of true DMCTs. Correspondingly, the specificity for this cell-type was defined as 1- FPR, where the false positive rate (FPR) is the ratio of false positives to the total number of true negatives. Of note, a true DMCT predicted to be a DMCT but where the predicted directionality of DNAm change is wrong, was counted as a false positive. Likewise, a true negative CpG predicted to be a DMCT (regardless of directionality) also defines a false positive. Finally, the precision or PPV for a given cell-type was defined as the fraction of correctly predicted DMCTs (with correct directionality of change) among all CpGs declared to be DMCTs.

*Real DNAm datasets with matched FACS cell counts:* We used the Illumina 450k dataset from Koestler et al [6] consisting of 18 samples, of which 6 were whole blood (WB) and 12 were experimentally reconstructed "whole blood" mixtures. This dataset is available from GEO under accession number GSE77797. In the case of the 6 whole blood samples, flow-cytometric cell count estimates for the 6 major blood cell subtypes were available. For the experimental mixtures, the mixing proportions were determined by the experimentalist and therefore known without error. DNAm data was normalized and processed as previously described [2]. In addition we analysed Illumina EPIC DNAm data for a total of 162 whole blood samples with matched FACS cell counts. The EPIC DNAm data was processed using minfi [7, 8] and BMIQ [8]. This dataset is available from NODE under accession number (project ID) OEP000651, i.e. from https://www.biosino.org/node/project/detail/OEP000651

*Real EWAS datasets of smoking:* We obtained Illumina 450k data from Liu et al [35], encompassing whole blood samples for 689 individuals. The raw data is available from GEO under accession number GSE42861. Normalized DNAm data was obtained from the authors and further adjusted for type-2 probe bias using BMIQ [31]. We also obtained Illumina 450k DNAm data from Hannum et al [6], encompassing whole blood samples for 656 samples. The raw and normalized data is available from GEO under accession number GSE40279. Normalized DNAm data was obtained from the authors and further adjusted for type-2 probe bias using BMIQ [31]. We also used Illumina 450k data from 790 buccal swabs collected as part of the MRC1946 birth cohort NSHD study, and which was previously normalized and analysed by us [9]. This data is only available by submitting data requests to; see full policy at http://www.nshd.mrc.ac.uk/data.aspx. Managed access is in place for this 73 year old study to ensure that use of the data are within the bounds of consent given previously by participants, and to safeguard any potential threat to anonymity since the participants are all born in the same week.

*Application of CellDMC and TCA to smoking EWAS datasets:* In the case of the Liu et al whole blood cohort, we ran CellDMC by adjusting for rheumatoid arthritis status, age and gender, and with smoking status (encoded as 0 for never-smokers, 1 for ex-smokers and 2 for current smokers) as the outcome of interest, and at a cell-type resolution level of 2 cell-types (myeloid and lymphoid). Total myeloid and lymphoid fractions per sample were estimated by running EpiDISH [2] for our 7 blood cell subtype reference and then separately adding the fractions within the lymphoid and myeloid compartments. In the case of Hannum et al whole blood cohort, we ran CellDMC by adjusting for age and plate, with smoking status as the outcome of interest (0=never-smokers, 1=ex-smokers, 2=current smokers), and at a cell-type resolution level of 2 cell-types (myeloid and lymphoid). In the case of Hannum, adjusting for plate has the advantage that it also adjusts for center and ethnic group, as these were distributed in a plate-specific manner. In the case of the buccal swab cohort, we estimated total epithelial, total lymphoid and total myeloid fractions using HEpiDISH [9]. We then ran CellDMC at a resolution of these 3 cell-types with smoking status (0=never-smokers, 1=ex-smokers, 2=current smokers) as the outcome of interest. In this cohort, no adjustment for age or gender is necessary because the buccal samples were all from women, and collected at the same age (53 years old). Beadchip and position effects were minor, and not deemed necessary to adjust for them, in line with our previous studies [10, 11].

TCA was run in two different ways, following the options offered by the authors of the TCA paper. One option, favoured by the authors, is to run TCA using the *tcareg* function. Pseudocode illustrating this mode is shown below:

```
# Build TCA model
tca.mdl <- tca(X = data, W = estFract, refit_W = F, C1 = cov.mod[,-1],
C2 = NULL,parallel = T,num_cores = 20)
# Use tcareg to get DMCTs
tcareg.o <- tcareg(X=data, tca.mdl=tca.mdl, y=smoking, C3 =
cov.mod[,-1], test = "marginal", null_model = NULL,
alternative_model = NULL, save_results = FALSE,
output = "TCA", sort_results = FALSE, parallel = T,
num_cores = 20, log_file = "TCA.log", features_metadata = NULL,
                debug = FALSE)
```

where *cov.mod* is the model matrix containing the same confounders as used in CellDMC (e.g. age). Thus, when running *tcareg,* we allow for potential confounding between outcome (smoking) and these factors through the *C3* argument (since the outcome is being used as the dependent variable). When applying the *tca* function, we also incorporate these factors through *C1* , so as to allow a cell-type specific effect of these confounders on the DNAm data. Association statistics are likelihood-ratio-test (LRT) statistics, from which P-values are derived.

In the second mode, we use the *tensor* function to infer the tensor object:

```
tca.mdl <- tca(X = data, W = estFract, refit_W = F, C1 = cov.mod[,-1],
C2 = NULL,parallel = T,num_cores = 20)
tca.tensor <- tensor(X = data, tca.mdl = tca.mdl)
```

and subsequently run separate "EWAS-regressions" for the inferred cell-type specific data-matrix. Thus, in this case we obtain t-statistics of association and corresponding P-values. We note that when running TCA in this second "tensor-mode", we did not adjust again for the confounders because we run the cell-type specific EWAS regressions which DNAm as the dependent variable, and therefore adjustment for the effect of confounders on DNAm was already done in the *tca*-step above.
